# Intelligence and Cognitive Flexibility Selectively Modulate Working Memory Functioning

**DOI:** 10.1101/2024.10.30.620994

**Authors:** Heming Zhang, Chun Meng, Xinyu Xu, Xin Hu, Ziqian Zhang, Benjamin Klugah-Brown, Bharat B. Biswal

**Author notes:** Correspondence: Chun Meng; Bharat B. Biswal.

## Abstract

Working memory is a key component of intelligence, which can be underpinned by task connectome, while cognitive flexibility plays a role in adaptive and executive function that supports working memory processes. However, it remains unknown how intelligence and cognitive flexibility modulate working memory functioning, in terms of behavioural performance and task-related brain hierarchy. Current study investigated the working memory task-related behaviour and brain connectomic gradient in healthy adults together with their intelligence and cognitive flexibility measures. Results showed cognitive flexibility and intelligence were not significantly correlated with each other but with working memory function. Principal component and partial least square analysis revealed that nearly half of variance was explained by their overlap component, which linked with task gradient values of frontoparietal and ventral attention network representing executive, control and cognitive functioning. Individuals with higher intelligence and cognitive flexibility showed less distance between frontoparietal/ventral attention network and unimodal regions in gradient axis during working memory performance. Further analyses demonstrated that task gradient of frontoparietal and ventral attention network was significantly associated with working memory function, and selectively mediated by cognitive flexibility and intelligence, which may suggest partly convergent neural mechanisms for higher order functions that are involved in working memory functioning. In conclusion, current study provides new insight into the relationship among task connectomic gradient, working memory, intelligence, and cognitive flexibility, which helps understand the multi-component model of working memory and underlying task gradient principle.

## 1. Introduction

Working memory is closely linked with intelligence and cognitive flexibility according to the widely accepted multi-component model (Baddeley, 2000; 2012). It shows that working memory process involves the central executive system, phonological loop, visuospatial sketchpad, and episodic buffer. Among these components, the phonological loop, visuospatial sketchpad, and episodic buffer are subsystems responsible for input and storage of information, controlled by the central executive system. These components contribute to fluid intelligence, which refers to an individual’s general intelligence (Baddeley, 2003; 2012; Cowan, 2008). Thus, working memory plays a vital role in cognitive processes related to general intelligence, such as reasoning, learning, and understanding (Colom et al., 2004; Conway et al., 2003; Thiele et al., 2022). On the other hand, the central executive system coordinates and controls the functions of the subsystems, performing advanced functions such as attention control and information processing. Cognitive flexibility so called switching is one of the most important subfunctions of the central executive system, which is a crucial function of working memory process (Miyake et al., 2000). Due to the limited capacity of working memory, individuals require to continuously switch the memory target in the working memory tasks (Baddeley, 2003; Brydges et al., 2018; Colom et al., 2004; Miyake et al., 2000). Cognitive flexibility involves the ability to shift focus between newly arrived and previously stored information (Angelopoulou and Drigas, 2021; Cowan, 2012). Brain disorders such as major depressive disorder, schizophrenia, and Alzheimer’s disease usually demonstrate such dysfunctions as reduced intelligence, weakened cognitive flexibility, and impaired working memory process (Braun et al., 2021; Parola et al., 2020; Vance & Winther, 2021). Therefore, we hypothesized that there might be potentially strong interactions between working memory, intelligence and cognitive flexibility.

The accumulating studies demonstrate that working memory might rely on widely distributed functional connectivity networks (Zhang et al., 2022; Braun et al., 2021) to support temporary storage and manipulation of a limited amount of information necessary for complex tasks (Lustig et al., 2001; Malinovitch et al., 2022; Barrouillet & Lecas, 1999; Prabhakaran et al., 1997). Executive function is an important function of working memory process which included frontoparietal network (FPN) and dorsal/ventral attention network (DAN/VAN). These networks could substrate high-level representations of primary functions, provide top-down signals, and guide the flow of activity across distributed brain networks (D’Esposito & Postle, 2015; Cocuzza et al., 2020). Recently, executive function has been shown to be associated with the development and hierarchical organization of brain networks along the principal gradient (Keller et al., 2023). Using whole-brain functional connectivity (FC) profiles, Margulies and colleagues developed a framework to identify functional gradients across the brain, reflecting the hierarchical organization of cognitive processing (Margulies et al., 2016). The principal gradient, ranging from unimodal to transmodal regions, offers a simple but integrative framework to delineate organizational structure for higher-order cognitive functions (Margulies et al., 2016; Murphy et al., 2018; Smallwood et al., 2021). While unimodal regions support primary visual and sensorimotor information, transmodal regions are involved in the information integration and comprehension from unimodal regions (Murray et al., 2014; Ito et al., 2020; Raut et al., 2020). The gradient difference between two brain regions or groups indicates their functional distance in the cortical hierarchical architecture or altered hierarchical organization in the human brain (Cross et al., 2021; Liang et al., 2021). Previous study reported the association between principal gradient of resting state brain connectome and task states (Ito et al., 2020; Wang et al., 2020), as well as individual cognitive ability (Huo et al., 2022). Our recent study found that the cognitive load during working memory task affected the principal gradient of task state brain connectome (Zhang et al., 2022). However, it remained unclear whether cognitive flexibility and intelligence affect principal gradient of brain connectome during working memory task.

Cognitive flexibility can be assessed by switch cost (Siqi-Liu & Egner, 2020). Working memory operation could incur switch cost (De Baene and Brass, 2011; Qiao et al., 2017), referred to the decrease of cognitive flexibility, which can be characterized by increased response time and decreased accuracy (Basak and Verhaeghen, 2011a; Monsell, 2003). Previous work showed cognitive flexibility decreases with the increase of working memory load (Draheim et al., 2016; Hester and Garavan, 2005; Lemire-Rodger et al., 2019). Working memory tasks have demonstrated high prediction accuracy for cognitive flexibility (Jiang et al., 2020), and individuals with better working memory performance also exhibit better cognitive flexibility (Pettigrew & Martin, 2016). On the other hand, working memory and intelligence are also intertwined (Conway et al., 2003), with the possible neural substrate in FPN (Cole et al., 2012; Kane et al., 2005; Jiang et al., 2020). Individuals who have higher intelligence scores demonstrated lower connectomic reconfiguration between working memory and resting state (Schultz & Cole, 2016; Thiele et al., 2022). Working memory tasks have demonstrated high prediction accuracy for intelligence (Greene et al., 2018; Jiang et al., 2020). Although both cognitive flexibility and intelligence plays a role in working memory, they are not highly correlated with each other (Friedman et al., 2006; 2011). Therefore, we hypothesized that the intelligence and cognitive flexibility had an overlap cognitive component to convergently modulate the working memory performance. Moreover, this overlap may be associated with task gradient underpinning working memory functioning.

In summary, current study aimed to explore whether and how intelligence and cognitive flexibility affect unimodal-transmodal working memory gradient. We employ publicly available data from openneuro.org, which includes working memory task, task-switching task, and Wechsler intelligence scale scores. First, we explore the relationship between working memory performance, intelligence scores, and cognitive flexibility from a behavioural perspective. We then assess the relationship between the working memory gradient, intelligence, and cognitive flexibility and explore the functional characteristics of the related networks using cognitive decoding of meta-analysis. Cognitive decoding of meta-analysis is aimed to find the functional characterization of global networks and to clarify the potential cognitive implications of these intelligence and cognitive flexibility-related regions based on the previous study (Huo et al., 2022; Xia et al., 2022a; 2022b). Finally, we refer to the working memory-related regions identified from cognitive decoding results to further analyse the relationship between working memory, intelligence, and cognitive flexibility. We hypothesized that intelligence and cognitive flexibility convergently modulate the behavioural performance and task gradient of working memory, resulting in overlapped neural substrate. In addition, the regions with gradient score associated with intelligence and cognitive flexibility mainly located within transmodal regions.

## 2. Materials and methods

### 2.1 Data and preprocessing

The working memory task including behavioural and fMRI data, together with intelligence and flexibility information, were obtained for 130 healthy participants from the UCLA Consortium for Neuropsychiatric Phenomics (CNP) dataset (https://openneuro.org/datasets/ds000030). The detailed demographic information has been reported in previous literature (Poldrack et al., 2016). For details, in working memory task, subjects were first shown several yellow circles positioned pseudorandomly around a central fixation crossing. Later, a single green circle shown on the screen. Subjects were required to indicate whether that circle was in the consistent position as one of the yellow circles had been. Half the trials were consistent, and half were inconsistent. Before the scanning, all subjects had been trained to ensure they were proficient with the tasks.

The fMRI preprocessing was performed for each subject and each run of working memory task and resting state, by using SPM12 (https://www.fil.ion.ucl.ac.uk/spm/) in the MATLAB 2018a environment. Briefly, the first 5 volumes were discarded. The remaining images were realigned to the first image of each run and then normalized into the standard MNI space using the EPI template. Out of the 130 subjects, 25 subjects had missing data, and 39 subjects were excluded according to the head motion criteria (maximal cumulative head motion of 2 mm or 2 degrees or mean frame-wise displacement of 0.2 mm, Zhang et al., 2022).

Further preprocessing was conducted for FC analysis, such as nuisance variable regression (including 24 head motion time courses, white matter and CSF signal) (Behzadi et al., 2007), linear detrend, and band-pass filtering of 0.01 – 0.1 Hz. For task-related FC analysis, we further regressed out the task with finite impulse response according to previous literature (Cole et al., 2019).

### 2.2 Behavioural and activation analysis

In the CNP data, Wechsler Adult Intelligence Scale score was calculated to represent for individual intelligence, the opposite value of the switch cost of task-switching task was used to represent individual cognitive flexibility, and working memory function (WM function) is indicated by accuracy of the working memory task. In task-switching task, subjects were presented images with different color and shape and required to respond to the image based on the task cue. The task switched between responding the color or the shape of the image. On 1/3 of trials the instructions switched. The switch cost was calculated by the reaction time (RT) of switch condition minus the RT of repeat condition in the task switching task. We assessed the pair-wise Pearson correlation between the working memory function, intelligence and cognitive flexibility in SPSS 21 (IBM, Armonk, NY). The working memory task activation was mapped in line with prior work (Zhang et al., 2023a). Significant clusters for task activation were identified using an initial threshold of p < 0.001 and the false discovery rate of p <0.05 for multiple comparison correction.

### 2.3 Task gradient analysis

The brain connectome i.e. whole-brain FC network was constructed for working memory task and resting state by using Yeo-400-atlas, which parcellates whole brain into 400 cortical regions related to 7 functional networks (Schaefer et al., 2018). Pearson correlation was computed between regional averaged time series, followed by Fisher’s z-transformation. In the end, individual 400 ×400 FC matrix was obtained for the task and rest respectively. Next, principal gradient of working memory task was computed following our prior work (Zhang et al., 2022). Specifically, top 10% of connections per ROI in the FC matrix were retained to obtain the normalized angle matrix. This matrix could capture the interregional similarity in connectivity profles (Margulies et al. 2016; Hong et al. 2019; Larivière et al. 2020). Then, we applied diffusion map embedding to capture the gradient components which explain the variance of connectivity pattern based on individual FC network. We finally generated a gradient template based on averaged FC matrix of all subjects and working memory task, as well as resting state, and used to align individual gradient maps via Procrustes rotations (Murphy et al. 2019). The principal gradient of functional brain network can reflect the hierarchical organization of cognitive processing from uni- to transmodal cortex (Margulies et al., 2016). The regional gradient score indicates the similarity and distance of regional FC with the rest of brain (Cross et al., 2021; Liang et al., 2021). Notably, the gradient difference between two brain regions or groups indicates their functional distance in the cortical hierarchical architecture or altered hierarchical organization in the human brain. The statistical significance threshold was set to p < 0.05.

### 2.4 Partial least squares analysis

Because intelligence and cognitive flexibility both correlated with task performance, we first performed principal component analysis (PCA) to obtain the overlap and orthogonal component between intelligence and cognitive flexibility. The overlap component may represent the convergent impact of intelligence and cognitive flexibility on working memory functioning. Next, the latent patterns of principal gradient related to working memory, overlap and orthogonal components of intelligence and cognitive flexibility, were investigated by using partial least squares (PLS) analysis (Figure 2A). PLS provides a multivariate approach to disentangle the latent relationships between brain and behaviour (Sun et al., 2022; Wei et al., 2022). It maximizes the covariance between brain and behaviour matrices by deriving the optimal linear combinations (i.e., latent components, LCs) of the original matrix (McIntosh and Lobaugh, 2004). In the current study, the gradient scores of Yeo-400 ROIs were stored in matrices X, and behaviour data were stored in matrices Y. Firstly, X and Y were normalized across all subjects, and the covariance matrix R was computed, followed by singular value decomposition of R into the singular vectors U and V, as well as the corresponding singular values. Statistical significance of the LCs was assessed using permutation testing (1000 permutation). Next, the composite scores L_X_ (brain score) and L_Y_ (behavioural score) were computed by projecting X and Y onto corresponding singular vectors V and U. The L_X_ and L_Y_ contain the weights such that the original variables (X and Y) maximally covary (Krishnan et al., 2011; Zeighami et al., 2019). To infer the contribution of the original variables (X and Y) to the LCs’ structure, brain loadings were defined by the Pearson correlation between X and L_X_. Similarly, behaviour loadings were defined by the Pearson correlation between Y and L_Y_. The stability of brain loadings was estimated by using 500 times of bootstrap tests and Bootstrap ratio (BSR) (Kebets et al., 2019). The significance threshold was set to p < 0.05 with FDR correction.

### 2.5 Cognitive decoding of meta-analysis

To find the functional characterization of global networks and to clarify the potential cognitive implications of these intelligence and cognitive flexibility-related regions, we conducted a meta-analysis based on the Neurosynth dataset (https://neurosynth.org/), following previous study (Huo et al., 2022; Xia et al., 2022a; 2022b). After obtaining the whole-brain correlation map between task gradient and the overlap as well as orthogonal component, Fisher’s r-to-z transformation was performed. The whole-brain correlation map was divided into positive correlation and negative correlation map respectively. Then, we obtained cognitive terms based on the Neurosynth dataset. The non-cognitive terms were discarded such as anatomical terms and brain network names. In the end the top 30 cognitive terms were reported in line with previous study (Xia et al., 2022a; 2022b). Permutation tests were performed to estimate the significance of correlation coefficient for each cognitive term. In brief, 10,000 random maps were constructed with smoothing and rescaling by using the variogram of real map. Thus, the random maps could preserve the spatial autocorrelation. Next, to be consistent with the real map, the first three networks with the highest loadings were chosen from the random maps, which were divided into positive correlation maps and negative correlation maps. Then, each random map was cognitively decoded in the Neurosynth database, and the results of 10,000 cognitive decoding were treated as the null model. The position of the actual correlation coefficient of each cognitive term in the null model was calculated as the p-value. Finally, the significance threshold was set to p < 0.05 with FDR correction.

### 2.6 Relationships between working memory, cognitive flexibility, and intelligence

According to PLS and cognitive decoding results, the somatomotor network (SOM), ventral attention network (VAN), and frontoparietal network (FPN) were significantly related to intelligence and cognitive flexibility, while the VAN and FPN were correlated with memory-related processes. So, VAN and FPN were utilized in further analyses. Firstly, we examined the working memory effect on the gradient of VAN and FPN, as well as the relationships between gradient of VAN/FPN and task performance. Secondly, linear mediation analyses were used to investigate whether intelligence and cognitive flexibility jointly or separately mediated the relationship between task gradient of VAN/FPN and working memory performance, by using SPSS (version 21, IBM, Armonk, NY). For example, we examined the relationship between the brain score and behaviour score (a path), the relationship between behaviour score and working memory function (b path), the direct effect of the brain score on working memory function after including behaviour (i.e. intelligence and cognitive flexibility) as a mediator in the model, and the indirect effect (c’ path, i.e., a*b path). The significance of the indirect effect was tested by using 5000 times bootstrapping and 95% confidence interval (CI) (Preacher and Hayes, 2008; Baum et al., 2017).

## 3. Results

### 3.1 Behavioural results and working memory task activation

Subjects of this study had intelligence (mean=28.73; SD=3.56), cognitive flexibility (mean=-0.15; SD=0.10) and working memory function (mean=0.89; SD=0.06). Significant positive correlation was found between flexibility and working memory function (r = 0.3728, p < 0.01; Figure 1B), as well as between intelligence and working memory function (r = 0.2500, p < 0.05; Figure 1B). This indicates that individuals with higher intelligence and cognitive flexibility performed better on the working memory task. However, the correlation between cognitive flexibility and intelligence was not significant (r = 0.0237, p > 0.05; Figure 1A). Using PCA, the orthogonal and overlap component were identified for cognitive flexibility and intelligence, resulting in positive correlation between the overlap component and working memory function (r = 0.4457, p < 0.05; Figure 1C). The orthogonal component was not correlated with working memory function (r = −0.086, p > 0.1; Figure 1C). Working memory task activation was found in dorsolateral prefrontal cortex, middle and inferior frontal gyrus, superior and inferior parietal gyrus, anterior cingulate cortex, supplementary motor area, and so on, which mainly aligned with VAN and FPN.

**Figure 1.**
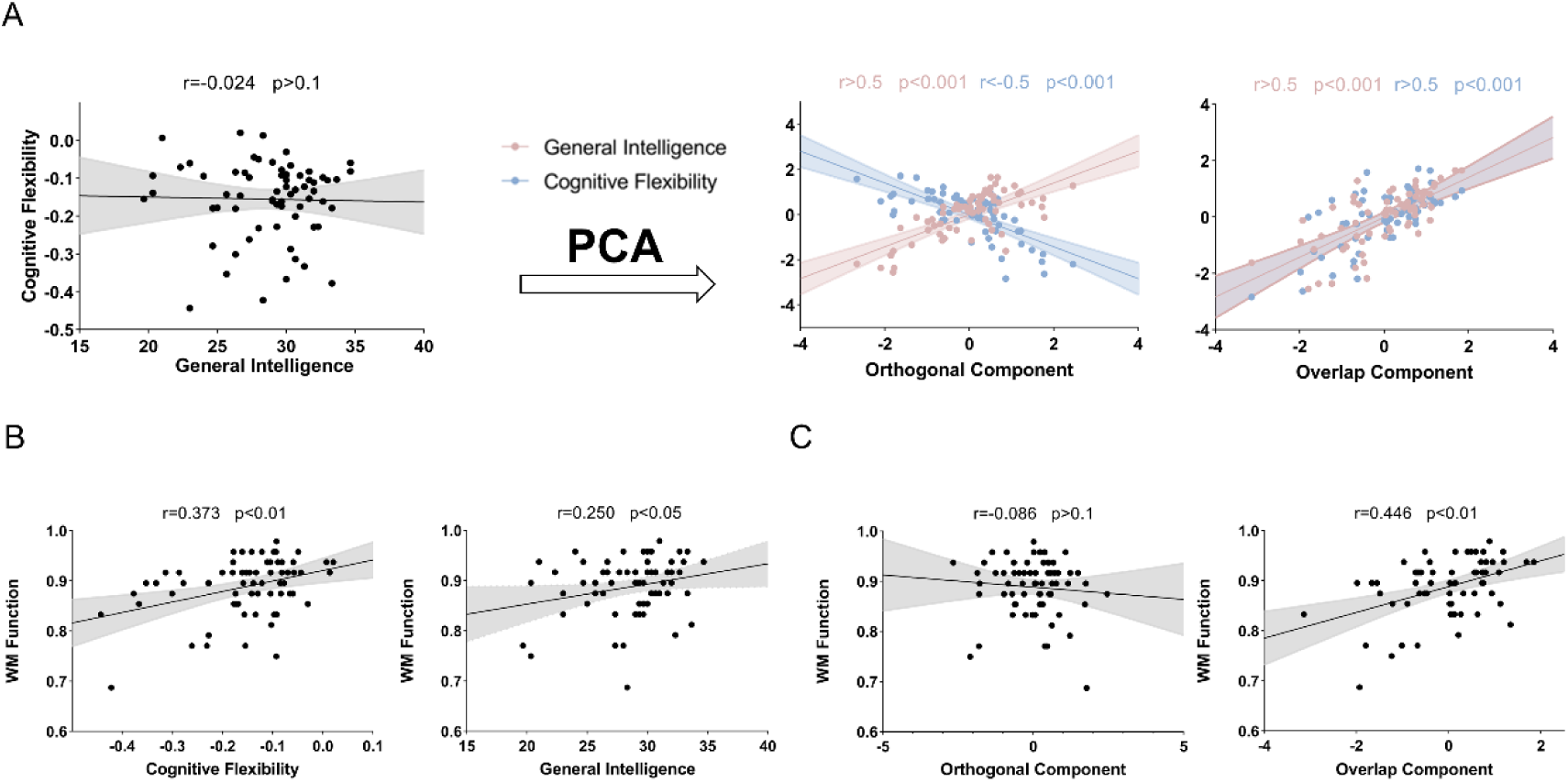
Behavioural results. (A) Correlation between general intelligence and cognitive flexibility was not significant. Using PCA, overlap and orthogonal components were identified, with comparable explained variance. (B) Both correlations between intelligence/ cognitive flexibility and working memory (WM) function were significant. (C) WM function was not significantly correlated with the orthogonal component, but with the overlap component.

### 3.2 Gradient pattern and association between working memory, cognitive flexibility, and intelligence

The principal gradient of working memory task averaged across all subjects showed a uni-to-transmodal pattern, with the SOM situated at the lowest end and the default mode network at the highest end (Figure 2B). This result is consistent with previous study, suggesting a stable hierarchical architecture (Zhang et al., 2022). The PLS results showed that task gradient of the SOM, VAN, and FPN were significantly correlated with the overlap component of cognitive flexibility and intelligence (Figure 2C; p < 0.05), but not with the orthogonal component of intelligence and cognitive flexibility. To further understand the functional characteristics of the global network in working memory functions, Neurosynth was used for cognitive decoding in the current study to identify cognitive functions related to the correlation maps (Yarkoni et al., 2011). The results showed that, for the positive correlation map, brain regions were significantly correlated with cognitive-related functions, such as cognitive control, executive function, and working memory (Figure 2D, FDR corrected p < 0.05). For the negative correlation map, the brain regions were significantly correlated with primary sensorimotor processes (such as tactile and motion) (Figure 2D, FDR corrected p < 0.05). Moreover, correlation between the brain score and working memory function was not significant (r = −0.12, p = 0.327; Figure 3A), but correlation between behavioural score and working memory function was significant (r = −0.42, p < 0.001; Figure 3B). Results of mediation analyses showed that the behavioural score mediated the association between brain score and working memory performance (Figure 3C).

**Figure 2.**
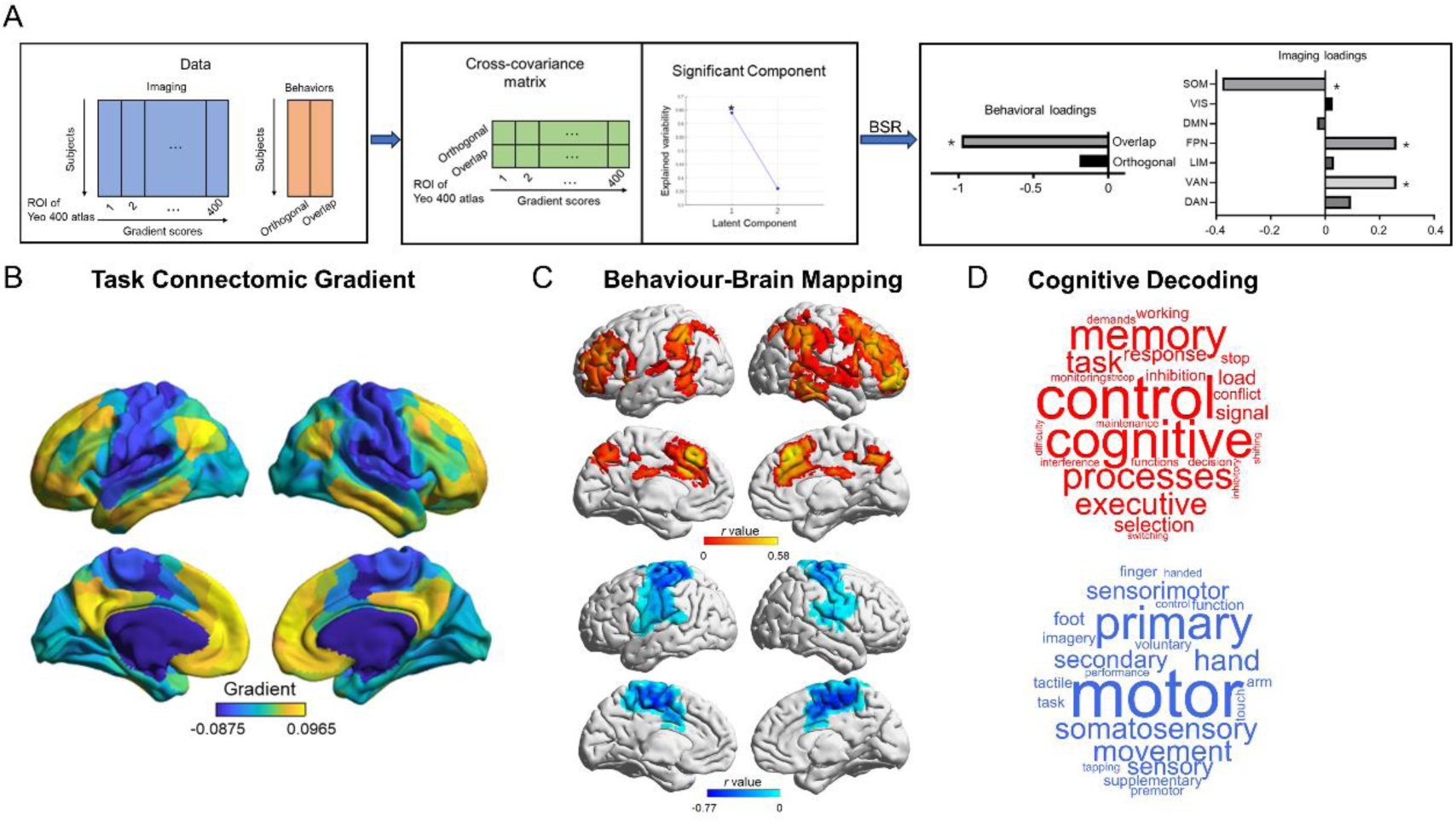
Principal gradient pattern based on working memory task state connectivity displayed pronounced correlation with the overlap component, which was related to brain regions and functions of working memory. (A) Processing pipeline of partial least squares (PLS) analysis, which identified significant latent components and multivariate loadings using the permutation testing and bootstrap ratio (BSR). (B) The average principal gradient pattern across all subjects demonstrated the whole brain topographic organization of task connectome during working memory task, including the lower end at SOM and VAN, as well as higher end at DMN and FPN along the gradient axis. (C) Brain regions that significantly contributed to the behaviour-brain association, as revealed by PLS, suggested that gradient pattern of SOM, FPN and VAN link with the overlap component of individual intelligence and cognitive flexibility. The red indicated the positive correlations while the blue indicated the negative correlations (FDR-corrected p < 0.05). (D) Cognition decoding analysis revealed the pronounced meta-analytic functions of significant clusters, based on Neurosynth.

**Figure 3.**
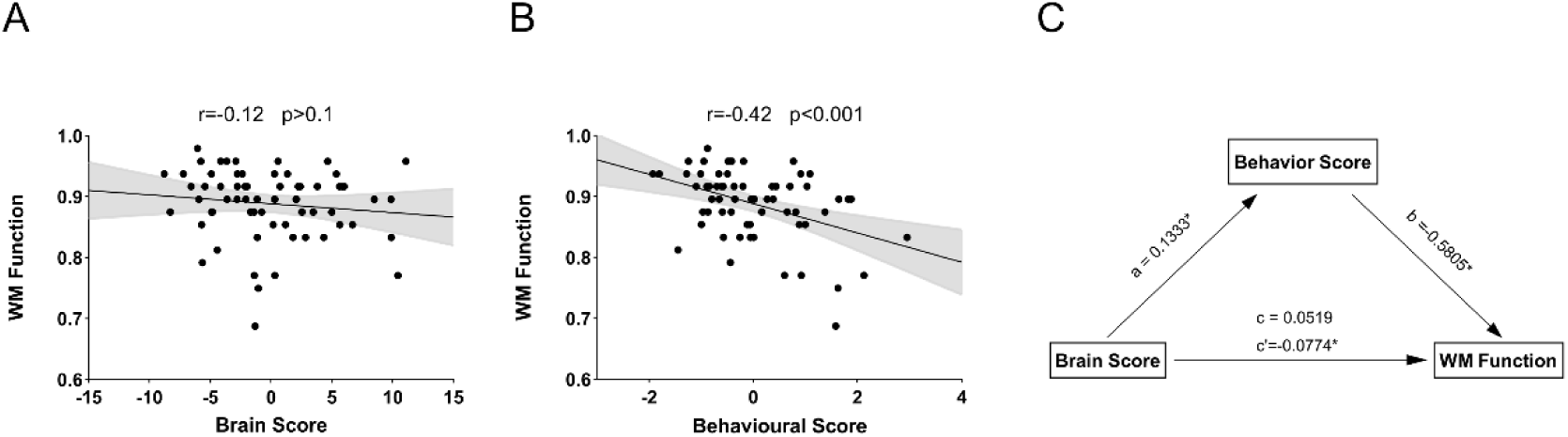
Results of correlations and mediation indicated that both brain and behavioural score could modulate individual working memory function. The brain score was not significantly correlated with working memory function (A), while the behavioural score was significantly correlated with working memory function (B). The behavioural score mediated the association between brain score and working memory function. Indirect effect (c’= – 0.0774) was assessed using bootstrapped confidence intervals [–0.1353 to –0.0249] (C). The asterisk (*) indicated p < 0.05.

### 3.3 Specific gradient patterns linked with FPN and VAN

Based on the results of the PLS method and cognitive decoding, we explored the gradient score of VAN and FPN which might interact with the overlap of intelligence, cognitive flexibility, and working memory functioning. There was a trend that the gradient score of VAN and FPN in working memory task was lower than in rest. The brain-behaviour correlation results showed that the gradient of VAN had a significant correlation with overlap of the intelligence and cognitive flexibility (r = −0.28, p < 0.05; Figure 4A). The gradient of FPN had no significant correlation with overlap component (r = −0.21, p = 0.093) but a significant correlation with intelligence (r = 0.31, p < 0.05; Figure 4C). Nonetheless, there was no significant correlation between the gradient score and cognitive flexibility (VAN: r = 0.24, p = 0.052; FPN: r = −0.02, p = 0.90), or the orthogonal component of cognitive flexibility and intelligence (VAN: r = 0.06, p = 0.63; FPN: r = −0.22, p = 0.07). Further mediation results demonstrated that the intelligence and cognitive flexibility selectively mediated the association between gradient score of the VAN/FPN and working memory function (Figure 4B, 4D).

**Figure 4.**
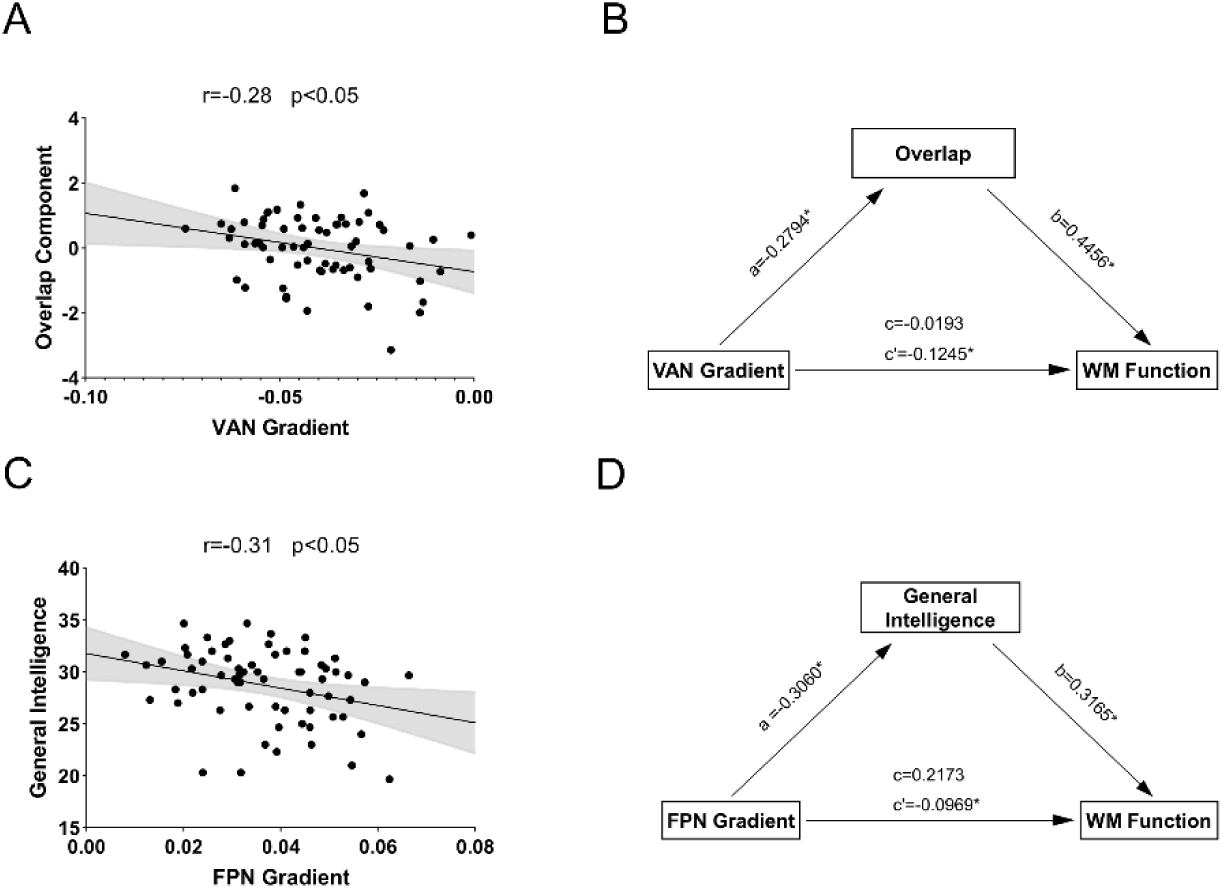
Mediation effect of intelligence and cognitive flexibility. (A) The overlap component was significantly correlated with average gradient score of VAN. (B) The overlap component, meaning both intelligence and cognitive flexibility, mediated the association between cortical hierarchy of VAN and working memory c. Significance of indirect effect (c’= –0.1245) was assessed using bootstrapped confidence intervals [–0.2476 to –0.0197]. (C) Only intelligence was significantly correlated with average gradient score of FPN. (D) Only intelligence mediated the association between cortical hierarchy of FPN and working memory function. Significance of indirect effect (c’= –0.0969) was assessed using bootstrapped confidence intervals [–0.2449 to –0.0040]. The asterisk (*) indicated p < 0.05.

## 4. Discussion

The current study investigated how intelligence and cognitive flexibility selectively modulate working memory performance using behavioural and cortical hierarchical analyses. The results showed that the overlap of intelligence and cognitive flexibility modulated working memory functioning, particularly mediated the relationship between cortical hierarchy of VAN and working memory function. Meanwhile, intelligence separately mediated the association between the cortical hierarchy of the FPN and working memory performance. Specifically, individuals with a less distance between VAN and unimodal regions in gradient axis were responding higher intelligence and cognitive flexibility, as well as better working memory performance. Moreover, individuals with a less distance between FPN and unimodal regions in gradient axis were responding higher intelligence as well as better working memory performance. Our findings confirmed the multi-component model of working memory, specifically both intelligence and cognitive flexibility jointly modulate working memory functioning.

The behavioural results showed that cognitive flexibility and intelligence had significant positive correlation with working memory function. These findings suggest that individuals with higher intelligence and cognitive flexibility tend to have better working memory performance. Previous study has shown that individual differences in working memory account for a significant amount of unique variance in intelligence (Shelton et al., 2010), and that working memory can predict intelligence (Oberauer et al., 2008). Additionally, recent study has highlighted the role of recoding information into a compact representation in working memory as a determining factor of intelligence (Chekaf et al., 2018). On the other hand, cognitive flexibility decreased with the increase of cognitive load (Draheim et al., 2016; Lemire-Rodger et al., 2019; Basak and Verhaeghen, 2011b), and subjects with better working memory performance corresponded to better cognitive flexibility (Pettigrew & Martin, 2016; Jiang et al., 2020). However, there was no significant correlation between cognitive flexibility and intelligence, which is consistent with previous studies showing no direct relationship between cognitive flexibility and intelligence (Friedman et al., 2006; 2011). Further analysis using PCA method showed that only the overlap of intelligence and cognitive flexibility was significantly related to working memory function, supporting the notion that intelligence and cognitive flexibility had similar cognitive function that modulate working memory performance (Herd et al., 2014).

The task gradient patterns were consistent with previous findings about cortical hierarchy of functional brain connectome during working memory task (Zhang et al., 2022). Using PLS analysis and cognitive decoding of meta-analysis, we characterized the functional networks associated with intelligence and cognitive flexibility effects. The results showed that the overlap of intelligence and cognitive flexibility was significantly correlated with the gradients of SOM, VAN, and FPN, while the orthogonal component of intelligence and cognitive flexibility was not related to the gradient of the whole brain network. These results were consistent with behaviour results, indicating that intelligence and cognitive flexibility can convergently modulate working memory behaviour and neural substrate (Herd et al., 2014). Brain areas included in VAN and FPN displayed positive correlation with the overlap component, which were associated with working memory, executive functions, and cognitive control by using cognitive decoding. The dorsolateral prefrontal cortex, anterior prefrontal cortex, inferior parietal cortex, and medial frontal cortex are usually implicated in memory load (Barch et al., 2013; Jimura et al., 2018; Lamichhane et al., 2020), while the lateral prefrontal cortex, premotor cortex, intraparietal sulcus, thalamus, and occipitotemporal regions are involved in working memory maintenance (Rypma et al., 1999; D’Esposito et al., 2000; Gazzaley et al., 2004). Cognitive flexibility is one of the subfunctions of the central executive system, which is an important function of working memory process (Miyake et al., 2000). The middle frontal regions involved attentional control during task switching (Mitsuhashi et al., 2022). The FPN contributes to domain-general task switching, while the premotor cortex contributes to perceptual switching (Gazes et al., 2012; Kim et al., 2012). This viewpoint is also supported by our result that regions negatively correlated with the overlap of intelligence and cognitive flexibility were the primary sensorimotor cortex, which is primarily involved in sensory and motor processes. Recent review has shown that the FPN and VAN are thought to be most important for the diverse forms of cognition that facilitate executive function (Bertolero et al., 2015; Rawls et al., 2022). They may undergo a relatively greater degree of integration as they mature, developing stronger functional connections with other systems and likely facilitating between-system communication in a manner that supports executive function (Baum et al., 2017; Bertolero et al., 2017; Cohen et al., 2016; Grayson & Fair, 2017; Oldham et al., 2022). These regions are associated with cognitive flexibility, visual abilities, and spatial abilities, which are important for general intelligence (Shi et al., 2022). The mediation analysis also supported that intelligence and cognitive flexibility jointly modulate working memory performance.

We further analysed the gradient scores of VAN and FPN. Intelligence scores exhibited significant negative correlations with the gradient score of the FPN in working memory task. The overlap of intelligence and cognitive flexibility had significant negative correlations with the gradient score of the VAN. These results consistently showed that individuals with higher intelligence and cognitive flexibility showed less distance between VAN/FPN and unimodal regions (such as SOM) in gradient axis during working memory performance. The FPN is important in diverse forms of cognition process (Jimura et al., 2018; Lamichhane et al., 2020). Our recent study found that the working memory task increased functional network integration and affected the principal gradient of task state brain connectome than resting state (Zhang et al., 2022; 2023b). Network integration has been shown to facilitate successful task performance in high cognitive tasks (Kitzbichler et al., 2011; Murphy et al., 2009; Zhang et al., 2022). The findings may imply that intelligence and cognitive flexibility requires sensomotor regions to cooperate with FPN in functional connectome (i.e. network integration) (Baum et al., 2017; Bertolero et al., 2017; Oldham et al., 2022). On the other hand, the SOM and VAN are both located at the lower end of the gradient axis, involved in sensomotor and attentional process (Margulies et al. 2016). The increased homogeneity within sensomotor regions and VAN may reflect the better local specialized processing in individuals with high cognitive ability (Huo et al., 2022). These results together indicate that individuals with higher intelligence and cognitive flexibility exhibit higher network integration in FPN and unimodal regions, as well as increased homogeneity within sensomotor regions and VAN, which in turn facilitates task performance (Kitzbichler et al., 2011; Murphy et al., 2009; Zhang et al., 2022).

Furthermore, our mediation results also supported that intelligence and cognitive flexibility jointly modulate working memory performance, with individual intelligence significantly mediating the association between the gradient of FPN and working memory function. Previous study has postulated a strong correlation between working memory and intelligence (Colom et al., 2004; Conway et al., 2003; Kane et al., 2005), as well as cognitive flexibility (Draheim et al., 2016; Hester & Garavan, 2005; Pettigrew & Martin, 2016). The current study brings together evidence to elucidate that hierarchical organization of FPN is a potentially effective way to predict the working memory performance. Besides, the intelligence and cognitive flexibility may play specific and necessary roles for improving working memory performance. Previous study confirmed multi-component model of working memory using behavioural or neuroimaging data and found that working memory is strong correlated with intelligence (Chekaf et al., 2018; Wongupparaj et al., 2015) and cognitive flexibility (Draheim et al., 2016; Lemire-Rodger et al., 2019). Current study using behavioural and cortical gradient analyses showed that functional gradient could reflect working memory function. The association between function distance of VAN/FPN and unimodal regions in gradient axis and working memory functioning is selectively mediated by cognitive flexibility and intelligence. These results directly link the unimodal-transmodal gradient to working memory and also provide new insight into the relationship among working memory, intelligence, and cognitive flexibility, which helps understand the underlying neural mechanism of working memory functioning.

One potential limitation is that the current study only explored the effects of cognitive flexibility and individual intelligence scores on working memory performance. Cognitive flexibility and intelligence are two single and specific measure and not enough to index complex cognitive constructs (Kane et al., 2002; Miyake et al., 2000). In other words, working memory process as one of the most important cognitive constructs, is hard to say it could be described clearly by using only two specific measures. Future studies should consider multifactors to measure cognitive constructs (such as inhibitory control) and analyse the combined effects of multifactors on working memory performance and brain networks. Furthermore, working memory could also influences cognitive flexibility by allowing individuals to hold and manipulate multiple pieces of information, which is critical for task switching and adapting to new strategies.

## 5. Conclusion

In conclusion, using behaviour and task connectomic gradient measures, current study has confirmed that both intelligence and cognitive flexibility are important factors in the human working memory functioning, and added new evidence for neural underpinning that intelligence and cognitive flexibility can modulate the gradient organization underlying working memory performance specifically in VAN and FPN. These novel findings contribute to a better understanding about the multi-component model of working memory and underlying task gradient principle.

## Author Contributions

Heming Zhang: Conceptualization, Methodology, Software, Analysis, Writing – original draft, Writing - review and editing. Chun Meng: Conceptualization, Supervision, Methodology, Writing - review and editing. Xinyu Xu, Xin Hu and Ziqian Zhang: Methodology, Analysis. Benjamin Klugah-Brown: Methodology, Writing - review and editing. Bharat B Biswal: Supervision, Writing - review and editing.

## Data and code Availability

The data used in current study is openly available in the UCLA Consortium for Neuropsychiatric Phenomics (CNP) dataset (https://openneuro.org/datasets/ds000030). The core code and toolboxe used in current study is already available on line (https://miplab.epfl.ch/index.php/software/PLS; and http://github.com/MICA-MNI/BrainSpace).

